# Genomic and transcriptomic analysis revealed new insights into the influence of key T6SS genes *hcp* and *vgrG* on drug resistance and interbacterial competition in *Klebsiella pneumoniae*

**DOI:** 10.1101/2023.05.16.540999

**Authors:** Wanzhen Li, Xiaolan Huang, Dan Li, Xiaofen Liu, Xiaoying Jiang, Xingchen Bian, Xin Li, Jing Zhang

## Abstract

Type VI secretion systems (T6SSs) act as a molecular weapon in interbacterial competition and play an important role in cell-cell interactions. Different species of bacteria use their T6SSs to perform a variety of functions according to ecological niche. Therefore, it is necessary to better understand the T6SS potential of *Klebsiella pneumoniae* (*K. pneumoniae*), a common clinical opportunistic pathogen. Here, we conducted a genomic analysis on the evolution, T6SS, virulence and antimicrobial resistance of 65 *K. pneumoniae* in patients with different infections. And we combined transcriptome analysis after knockout of key gene in T6SS of this species. Results showed that genes encoding a T6SS were present in all *K. pneumoniae* in this study, and there was no correlation was found between T6SS cluster and carbapenem resistance and virulence genes. Differentially expressed genes (DEGs) including 1298 co-upregulated and 1714 co-downregulated were identified after *hcp* or *vgrG* deletion. Kyoto Encyclopedia of Genes and Genomes (KEGG) pathways analysis have demonstrated common changes in quorum sensing, propionate metabolism and other pathways. And we found that the deletion of *hcp* or *vgrG* genes up-regulated of beta-lactam (*bla*_KPC-2_) and other resistance genes. Interbacterial competition experiments showed that *hcp* and *vgrG* are essential genes for competitive ability of ST11 *K. pneumoniae*. Taken together, the entire study provides further insight into the investigation of T6SS in *K. pneumoniae* through genomic and transcriptomic analysis.

**Importance:** Gram-negative bacteria use T6SS to deliver toxin effectors to interact with neighboring cells for niche advantage. *K. pneumoniae* is an opportunistic nosocomial pathogen that often carriers multiple cope T6SSs, but the function of its T6SS has not yet elucidated. Here, we performed a genomic analysis of 65 clinical *K. pneumoniae* strains, in order to explore the relationship between T6SS and virulence and resistance genes. We also study the repertoire after knockout of key gene in T6SS of this species by transcriptomics. It was suggested that T6SS is associated with drug resistance, and its key genes *hcp* and *vgrG* are critical for the interspecies competition of *K. pneumoniae*.

## Introduction

The Type VI secretion system (T6SSs) is a nanomachine that is widely distributed in Gram-negative bacteria and can deliver toxin effectors into prokaryotic or eukaryotic cells in order to attain dominance within a given niche (1, 2). T6SSs is a phage-like device anchored to the bacteria membrane and usually consists of 13 conserved core components, namely TssA-M (3, 4). Among them, Hcp, VgrG, and PAAR are important T6SS structural proteins involved in the transport of effector proteins (5–7). The function of T6SS is mainly interbacterial antagonism, involving interspecies and intraspecies competition(8, 9). T6SS effector proteins with antibacterial activity have been reported to have the following functions: lysing essential macromolecular substances such as DNA, phospholipids and peptidoglycans of bacteria (9–14). In recent years, the functions of T6SS effector proteins have been further expanded, including against fungi and eukaryotic cells. For example, bacteria can manipulate T6SS to participate in host cells physiological processes, including adhesion, skeletal rearrangement and evasion innate immunity to gain additional advantages for colonization, dissemination and survival (15). In addition, T6SS can also perform mental acquisition (10, 16). In brief, T6SS has diverse functions and thus plays an important role in microbial adaptive survival.

*Klebsiella pneumoniae*(*K. pneumoniae*) is an important opportunistic Gram-negative pathogen which exists in the environment and can cause severe nosocomial infections in immune-compromised individuals (17). Multiple copies of T6SS are often present in *K. pneumoniae*, and most carrying two T6SS clusters ^[6]^. The prevalent carbapenem-resistant *K. pneumoniae* (CRKP) in China is ST11 type, which also carries two T6SS gene clusters (18). It has been reported that *K. pneumoniae* T6SS is contributed to bacterial competition, cell Invasion and colonization (19). Although there have been several studies on T6SS of *K. pneumoniae*, there are still many gaps in the ecology and pathogenesis of *K. pneumoniae* that need to be further explored.

In general, reports on T6SS of *K. pneumoniae* displayed a deficiency of definite clinical and molecular information. Genomic and transcriptomic analysis of Chinese *K. pneumoniae* strains is lacking. This study aimed to comprehensively understand the biological roles of T6SS in *K. pneumoniae*. We conducted a genomic analysis on the evolution, T6SS, virulence and antimicrobial resistance through 65 clinical isolates collected from patients with different infections. Then we focused on ST11 *K. pneumoniae* and combined transcriptome analysis after deletion of key gene *hcp* and *vgrG* in T6SS of this species.

## Results

### Identification of Type VI Secretion system in *K. pneumoniae* clinical isolates

The phylogeny of 65 *K. pneumoniae* clinical isolates was generated (Fig.1). Most strains (52 of 65) were isolated from patients with bloodstream infection. In ST11 *K. pneumoniae*, all but one of the 16 strains encoded the capsular polysaccharides (CPS) loci of KL64. Most of ST15 were KL19 (15 of 17), and all of ST23 were KL1. Most of the carbapenem resistant strains carried *bla*_KPC-2_ gene (29 of 30). These results showed that all clinical *K. pneumoniae* in this study carried the T6SS gene cluster, regardless of whether it contained carbapenem resistance or virulence genes.

**FIG 1.**
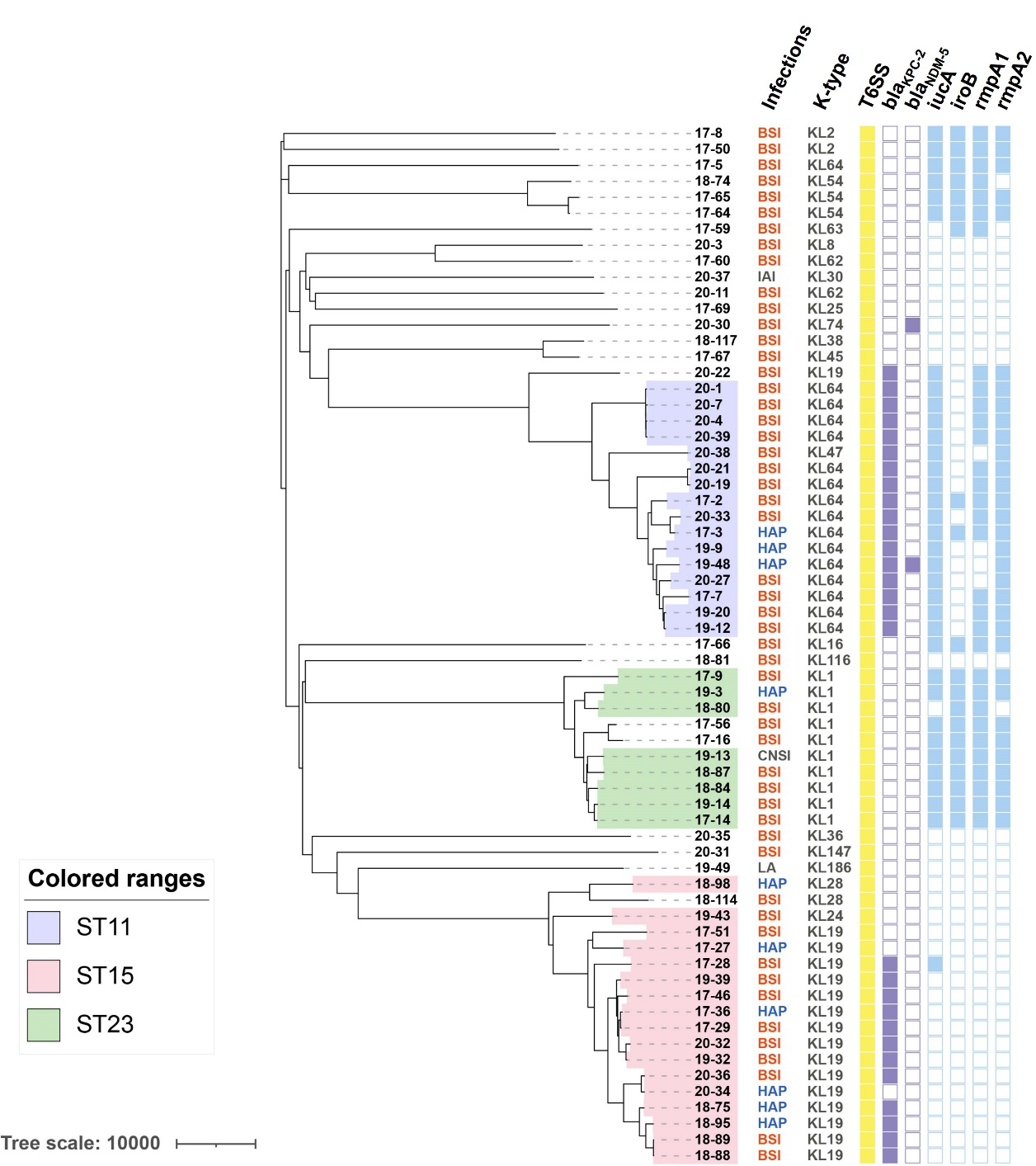
Phylogenetic tree of 65 *K. pneumoniae* strains. Different colored ranges represent different strains of ST type. Column, from 1 to 2, represents: column 1, type of infection; column 2, The CPS loci of the strain. Column, from 3 to 9, the color of square represents the exist of the indicated gene. BSI: bloodstream infection; IAI: intra-abdominal infection; HAP: hospital acquired pneumonia; CNSI: central nervous system infection; LA: liver abscess.

### Deletion of *hcp* or *vgrG* leads to changes in multiple physiological pathways of ST11 *K. pneumoniae* HS11286

To further investigate whether this T6SS is involved in other physiological pathways in *K. pneumoniae*, we deleted *hcp* or *vgrG* in ST11 *K. pneumoniae* HS11286 and conducted RNA-sequencing (RNA-seq) to compare its transcriptomic profiles with the wild-type strain (WT). The number of up- and down-regulated differential expression genes (DEGs) is shown in Fig. 2A. The total number of DEGs was similar in Δ*hcp* and Δ*vgrG* mutants (3329 *vs*. 3290 total DEGs, respectively). And there were 1298 co-upregulated genes and 1714 co-downregulated genes (|log2FoldChange| > 1, P-value<0.05) in WT/Δ*hcp* and WT/Δ*vgrG* comparisons. The top 10 Gene ontology (GO) analysis with the smallest P-values were selected for display (Fig. 2B). GO analysis showed that the both mutants of EDGs were partial similar in terms of biological process (BP) and cellular component (CC). However, The DEGs in both mutants were consistent only in transporter activity of molecular function (MF).

**FIG 2.**
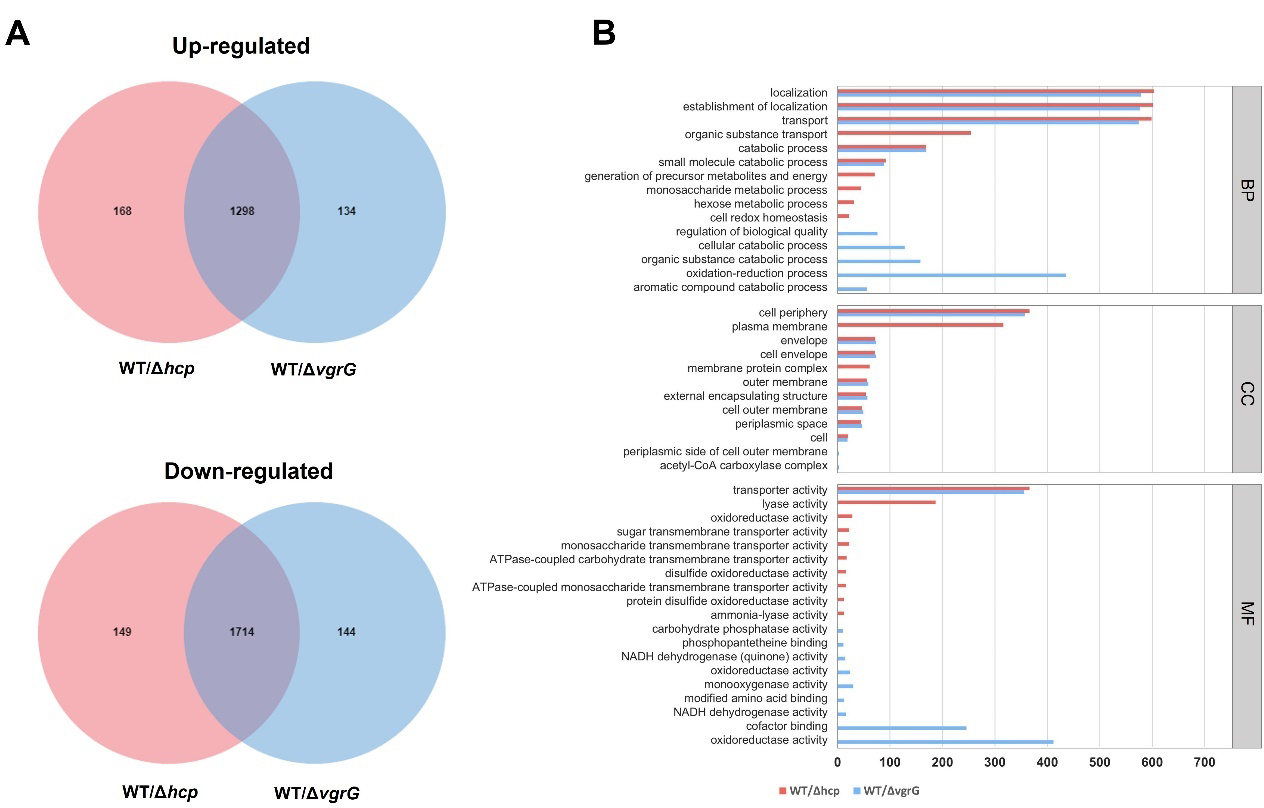
Comparative analyses of the transcriptional response to Δ*hcp* and Δ*vgrG K. pneumoniae* mutants. (A) Venn diagrams showing the significantly increased or decreased transcript abundance (|log2FoldChange| > 1, P-value&lt;0.05) in response to the different *K. pneumoniae* mutants. (B) GO analysis of up- and down-regulated DEGs in WT/Δ*hcp* and WT/Δ*vgrG*. BP: biological process; CC: cellular component; MF: molecular function.

The top 15 KEGG pathways with the smallest false discovery rate (FDR) values were selected for display, and the results are shown in Fig. 3. KEGG enrichment analysis revealed that 10 pathways overlapping after *hcp* or *vgrG* gene knockdown. There is quorum sensing, propanoate metabolism, phosphonate and phosphinate metabolism, oxidative phosphorylation, methane metabolism, etc. Notably, deletion of the *hcp* gene also affects the biofilm synthesis pathway, which is closely related to antimicrobial resistance. While *vgrG* gene deletion leads to altered fatty acid synthesis. Fatty acid synthesis can provide raw materials for bacterial cytosolic phospholipid synthesis, and this result suggests that *vgrG* may affect cell membrane perturbation.

**FIG 3.**
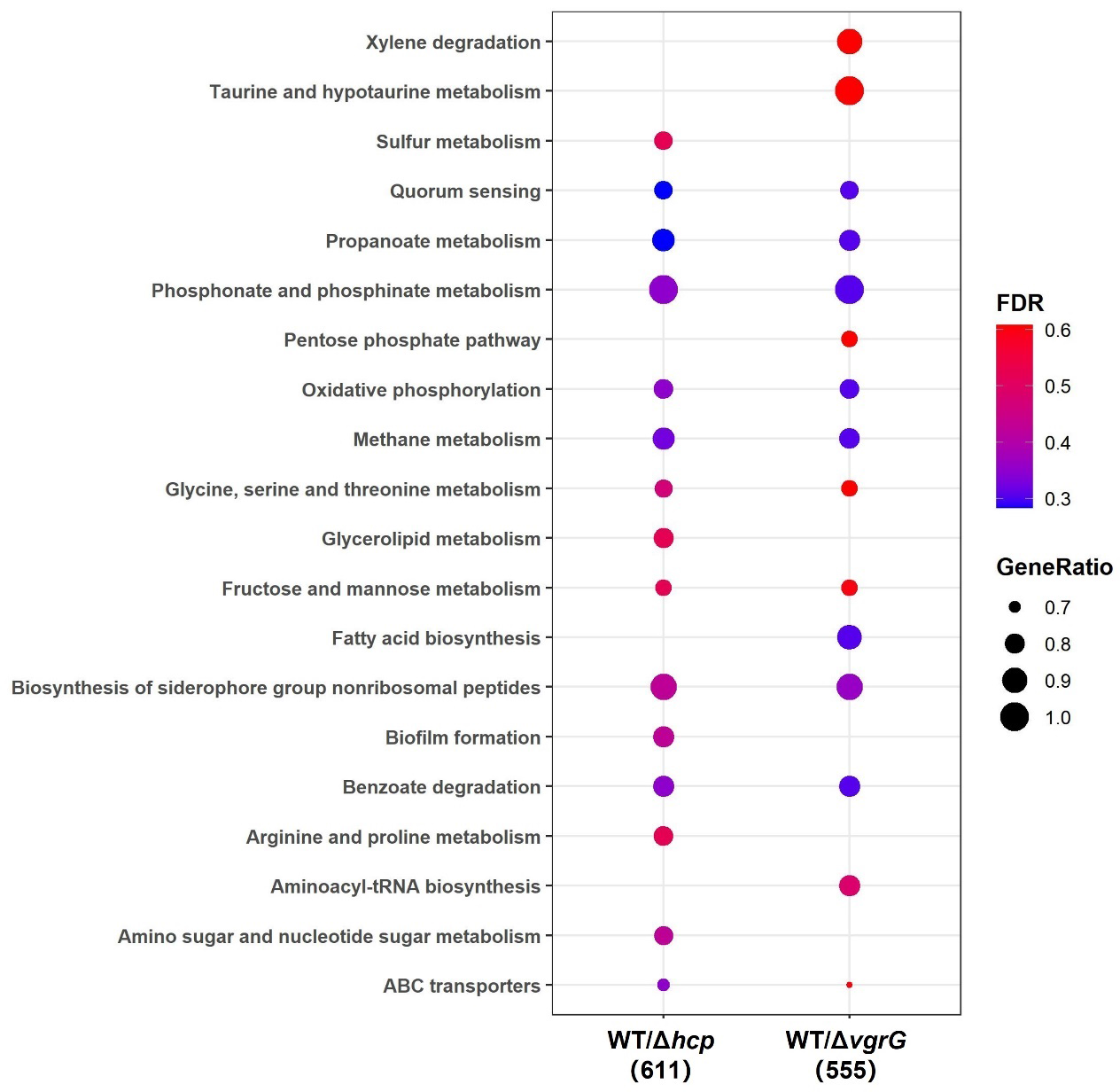
KEGG pathway enrichment of DEGs in the different *K. pneumoniae* mutants. GeneRatio represents the ratio of the number of DEGs enriched in one pathway to the number of all annotated DEGs.

### Deletion of *hcp* or *vgrG* up-regulated multiple resistance genes in HS11286

According to the DEGs, it was found that the deletion of *hcp* or *vgrG* would lead to the changes in drug resistance gene expression (Fig. 4). Both mutants caused up-regulation of resistance genes involved in aminoglycosides (*aphE*, *baeR*), beta-lactam (*bla*_KPC-2_*, bla*_TEM-1_), fluoroquinolone (*gyrA*, *parC*, *marR*, *marA*), macrolide (*mdtJ*, *mdtI*, *emrA*, *emrB*) and other resistance genes. However, *K. pneumoniae* HS11286 is a carbapenem resistant strain, so the change of resistance phenotype of the two mutant strains could not be observed from antimicrobial susceptibility test (see TABLE S1 in the supplemental material).

**FIG 4.**
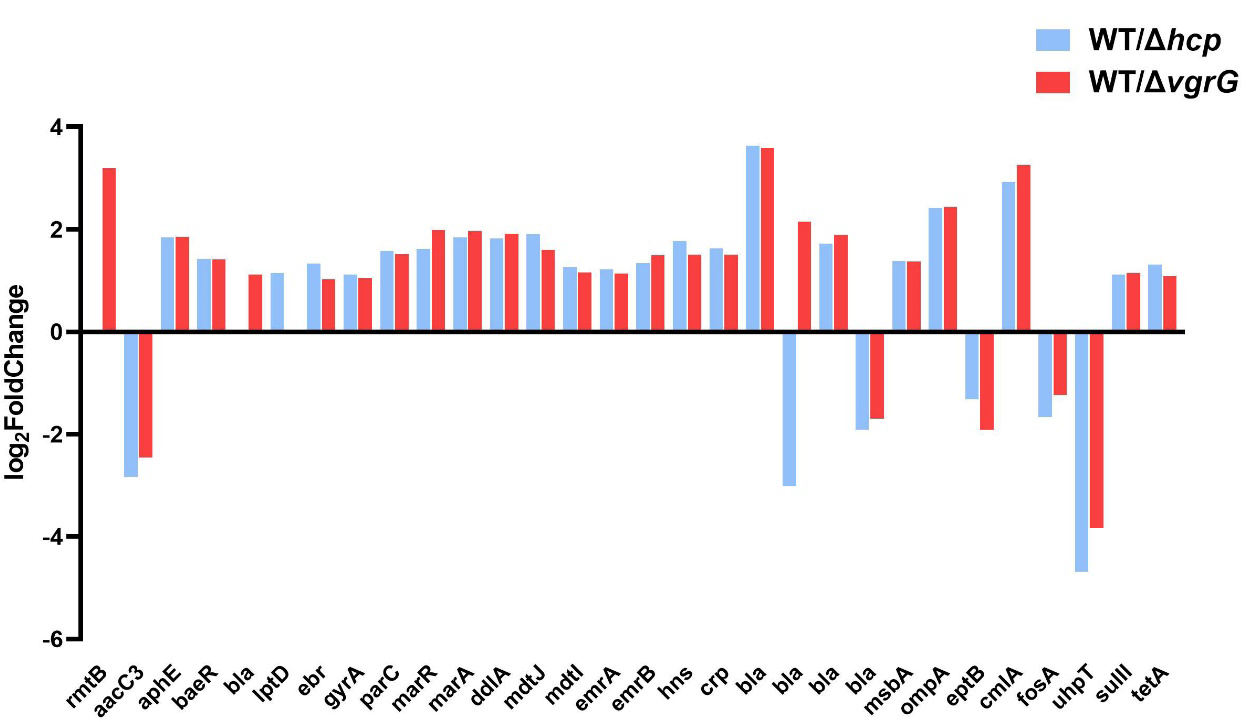
Fold changes in drug resistance genes after *hcp* or *vgrG* deletion.

### Transcriptional profile of T6SS-related genes in Δ*hcp* and Δ*vgrG* mutants compared to WT strains

*K. pneumoniae* HS11286 encodes two T6SS clusters (Fig. 5A), and locus I is the intact T6SS cluster with all conserved core genes. RNA-seq results revealed that deletion the *hcp* or *vgrG* genes of locus I can repress almost all T6SS-related genes of locus II (Fig. 5B). Further analysis revealed that after *hcp* gene knockdown, all T6SS-related genes in the HS11286 incomplete T6SS cluster were down-regulated, except for the *paar* gene. In the intact T6SS gene cluster, only *tssC*, *tssK*, *tssL* and *tagL* were down-regulated, while the rest of genes were not significantly changed (see TABLE S2 in the supplemental material). In the Δ*vgrG* mutant, all of the complement T6SS genes were down-regulated, while in the intact T6SS gene cluster, *tssC*, *tssK*, *tssL*, *tagL*, *clpV*, *tssM*, *tssF*, *tssG*, and *tssJ* were down-regulated. In addition, the immunity protein of T6SS locus I were down-regulated after *vgrG* deletion.

**FIG 5.**
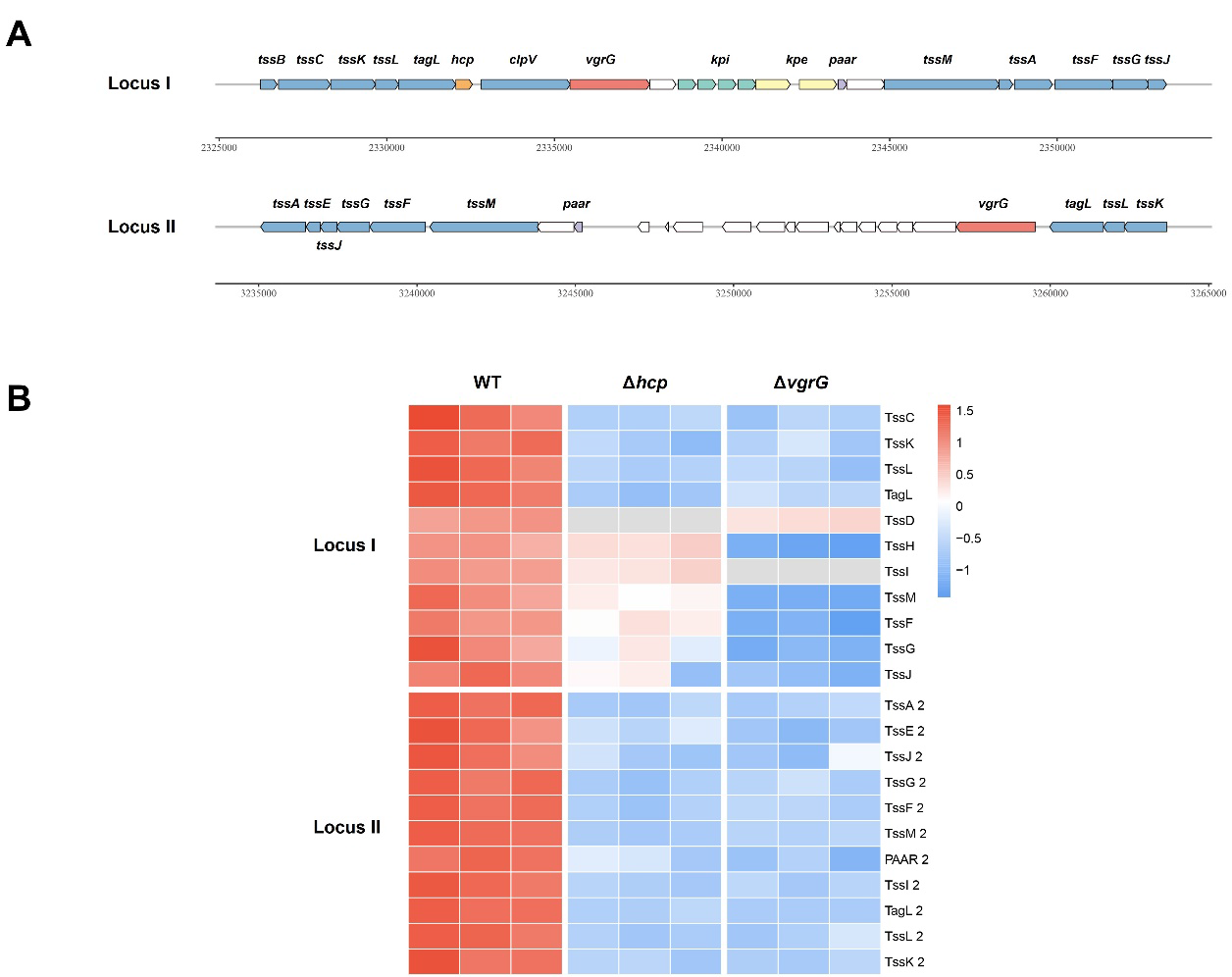
Differential expression of T6SS clusters in Δ*hcp* and Δ*vgrG K. pneumoniae* mutants. (A) Schematic representation of T6SS locus in *K. pneumoniae* HS11286. (B) Heatmap of Δ*hcp* and Δ*vgrG* mutants compared to WT strains (HS11286).

As observed in the RT-qPCR analyses (Fig S1). The expression levels of *clpV* in Δ*vgrG* was much lower (P < 0.05) than WT. The other two genes *hcp* and *paar* did not change significantly. All these results are consistent with the above transcriptome results. However, the *vgrG* and *clpV* expression of Δ*hcp* was higher than WT (P < 0.05), which was not consistent with the transcriptome results.

### Hcp and VgrG are required for interbacterial competitions in *K. pneumoniae*

To investigate the contributions of the two structural proteins Hcp and VgrG protein on the function of *K. pneumoniae* T6SS, we tested whether mutants exert antibacterial activity against *Escherichia coli* EC600 in a T6SS-dependent manner. Growth of EC600 was significantly reduced (2-fold) in the presence of WT when compared with the Δ*hcp* and Δ*vgrG* mutant strains (Fig. 6). The antibacterial ability of the Δ*hcp* and Δ*vgrG* mutant strain against EC600 were restored by complementation with plasmid containing the *hcp* or *vgrG* gene. Colony counting revealed that after 5 h of co-culture with EC600, the Δ*hcp* and Δ*vgrG* mutants reduced in vitro competitive ability compared to the WT, but there was no significant difference after statistical analysis (see Fig. S2 In the supplemental material). After 24 h co-culture with EC600, the survival rate of EC600 increased in the mutants group compared to the WT, and there was a statistically significant difference.

**FIG 6.**
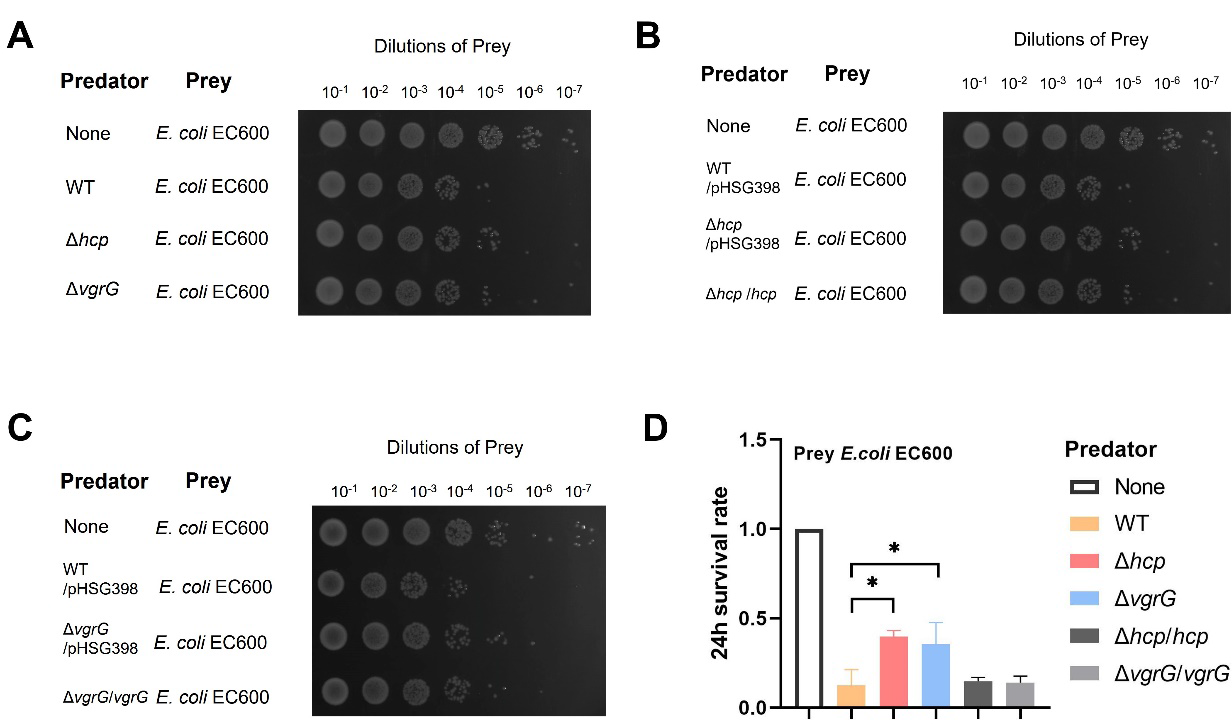
Interspecies killing by *K. pneumoniae* HS11286 in a contact-dependent manner. (A–C) Interbacterial competitive growth assays. Surviving *Escherichia coli* EC600 after 24 hours of coincubation with *K. pneumoniae* HS11286 (WT), Δ*hcp* mutants (Δ*hcp*), Δ*vgrG* mutants (Δ*vgrG*), WT containing the empty vector (WT/pHSG398), Δ*hcp* mutant containing the empty vector (Δ*hcp* /pHSG398), Δ*hcp* mutant complemented with pHSG398-*hcp* (Δ*hcp*/*hcp*), Δ*vgrG* mutant harboring pHSG398 (ΔvgrG/pHSG398), Δ*vgrG* mutant harboring pHSG398-*hcp* (Δ*vgrG*/*vgrG*). Recovered mixtures were plated onto Luria-Bertani medium supplemented with 100 μg/mL rifampicin. (D) Survival rates of the prey strain *Escherichia coli* EC600, after 24 hours of coincubation with the predator strains *K. pneumoniae* HS11286 (WT), Δ*hcp* mutants, Δ*vgrG* mutants, *hcp* and *vgrG* complemented strains. The data represent the means of 3 independent trials. \**P* &lt;0.05 by 1-way analysis of variance (Δ*hcp* mutants or Δ*vgrG* mutants compared with the wild-type strain).

## Discussion

Type VI secretion system plays a key role in remodeling bacteria due to its intraspecies and interspecies antibacterial activity (20–23). Based on the variation between different regions and different infections. We studied the distribution of T6SS in *K. pneumoniae* isolated from bloodstream, pneumonia and other infections. The second-generation sequencing results showed that each clinical isolated strain had T6SS gene cluster, and the identification of T6SS by PCR of key genes was limited. Moreover, PCR identification could not identify the number of copies of T6SS in the strain, and the existing method could not identify the copy number of T6SS in the second-generation sequencing results. It was found that no correlation between T6SS gene cluster and carbapenem resistance and virulence genes. However, the relationship between actual T6SS activity and drug resistance and strain virulence remains unclear.

Different bacteria use and adapt their T6SS apparatus to secrete effector proteins for specific functions, depending on the environmental niche of the bacteria (20, 24). The two T6SS structural proteins, Hcp and VgrG, have dual functions as both components and substrates of T6SS, and shed from the extracellular to the environment when T6SS is activated (25, 26). In addition, Hcp and VgrG are also the "carriers" of effector protein secretion(6). In this study, we found that the deletion of *hcp* or *vgrG* genes in T6SS gene cluster I can cause a variety of similar physiological pathway changes of the *K. pneumoniae* HS11286. At the same time, the lack of *hcp* or vgrG gene also caused the up-regulation of multiple resistance genes. Previous studies have shown that large conjugative multidrug resistance plasmids can repress the T6SS activity in *Acinetobacter baumannii* (27). It has also been reported that sub-inhibitory concentrations of β-lactam antibiotics can increase the activity of HS11286 T6SS and enhance the T6SS-dependent killing (13). Thus, the relationship between T6SS activity and drug resistance in *K. pneumoniae* needs further investigation.

Transcriptomic analysis showed that the deletion of *hcp* and *vgrG* genes also inhibited the expression of incomplete T6SS gene cluster. Whether the deletion of the key gene in T6SS will affect another T6SS gene cluster has not been studied in the previous literature, which needs to be further clarified.

Some strains carry multiple different copies of T6SS, suggesting a diversity of functions (28). *K. pneumoniae* also carries multiple copies of T6SS, for example, the current domestic ST11 usually carries two T6SS gene clusters. After the deletion of key genes *hcp* and *vgrG*, the expression of some genes in the complete T6SS gene cluster was also changed. Considering that the secretion process of T6SS is a cyclic process in which Hcp and VgrG proteins are injected into the extracellular before reassembling the components (25), it remains to be clarified whether the deletion of key proteins affects the assembly and secretion process. T6SS is generally considered to be a contact-dependent secretory system, these results showed that the knockout of *hcp* or *vgrG* genes affect the ability of T6SS to compete interspecific. Since T6SS is a contact-dependent bactericidal mechanism, these results also suggest that longer contact between strains, implying a stressful inter-strain ecological environment, would allow *K. pneumoniae* to exert its T6SS competitive advantage. This does not mean that T6SS is directly involved in the pathogenesis of *K. pneumoniae*. Instead, it is possible that T6SS can eliminate potential microbial competitors to benefit its own colonization. For example, *Yersinia pseudotuberculosis* uses T6SS-3 to secret nuclease effector protein Tce1 to kill other bacteria and facilitate gut colonization in mice (29).

In conclusion, T6SS was present in *K. pneumoniae* isolates of both bloodstream infection and pneumonia, and no association was found with carbapenem resistance. In CRKP strain HS11286, *hcp* and *vgrG* are essential proteins for T6SS activity of *K. pneumoniae*. Deletion of *hcp* and *vgrG* can cause similar physiological pathway changes and lead to up-regulation of many drug resistance genes in *K. pneumoniae*. Meanwhile, deletion of *hcp* or *vgrG* inhibited the expression of the locus II T6SS gene cluster in the strain. In conclusion, This study further expands the understanding of T6SS in *K. pneumoniae*.

## Materials and Methods

### Bacterial strains and plasmids

Bacterial strains, plasmids and primers used in this study are listed in TABLE S3-4. Sixty-five clinical isolates used in this study were kindly provided by Institute of Antibiotics, Huashan Hospital. The whole genome of these strains was sequenced using Illumina platform. *Kleborate* was used to identify the sequence type, K_locus, drug resistance genes and virulence genes of these strains (http://github.com/katholt/Kleborate) (30). Carbapenem-resistant *K. pneumonia* HS11286 (NC_016845) was isolated from clinical sputum specimen and belong to ST11 (31). *Escherichia coli* EC600 is the storage strain of the laboratory.

### Antimicrobial susceptibility test

Microdilution method was performed to determine the minimum inhibitory concentration (MIC) of each strain. The results were interpreted according to the Clinical and Laboratory Standards Institute (CLSI) (32).

### Gene deletion and complementation

The gene deletion strains HS11286*-* Δ*hcp* and HS11286*-*Δ*vgrG* were constructed using the λ Red recombinase method (33). For complementation, the DNA sequence of *hcp* and *vgrG* were PCR amplified from HS11286 and cloned into the pHSG398 plasmid (laboratory modification with apramycin resistance). These cloned plasmids were transferred into their corresponding gene deletion mutants by electroporation.

### Interbacterial competitive assays

Using the rifampin-resistant *Escherichia coli* EC600 as the prey strains. The indicated predator and prey strains were grown overnight. Cultures were diluted 1:100 with fresh LB and grown to mid-exponential phase, and resuspended in PBS to OD_600_ = 0.5 after centrifugation. Then mixed predator and prey strains at 10:1 ratio. The mixtures were spotted (10 μL) on LB agar with rifampin (100 μg/L), and incubated for 5 h or 24 h at 37°C. Harvested the mixture spots and plate serial dilutions on rifampin (100μg/L) selective agar.

### RNA-Seq and bioinformatic analyses

*K. pneumoniae* HS11286, HS11286*-Δhcp* and HS11286*-ΔvgrG* mutant strains were incubated overnight (37°C, 180 rpm). Overnight cultures were diluted 1:100 with fresh LB and grown to logarithmic phase (OD_600_ = 0.4-0.6), then collected. RNA extraction and sequencing was performed by Personalbio (Shanghai) using the Illumina Novaseq 6000 platform. Quality assessment and filtration of the raw data were performed by and Cutadapt. The filtered RNA-Seq reads were mapped to the reference genome (NC_016845) using Bowtie2 (34). Transcript abundance was quantified using fragments per kilobase per million mapped fragments (FPKM) normalization method. We use the DESeq2 (35) for differentially expressed genes (DEGs) analysis, conditions as follows: |log2FoldChange | > 1, P-value< 0.05. Venn diagrams were drawn by the online tool jvenn (http://jvenn.toulouse.inra.fr/app/example.html) (36).

We performed Gene Ontology (GO) and Kyoto Encyclopedia of Genes and Genomes (KEGG) functional enrichment analysis to identify which DEGs were significantly enriched in GO terms or metabolic pathways. GO terms and KEGG pathway with false discovery rate (q-value) < 0.05 were considered as significantly altered. According to the KEGG enrichment analysis results of DEGs, select the top 15 pathways with the smallest p-value and the most significant enrichment for display. All the enrichment visualization was performed using R.

### Quantitative real-time reverse-transcription polymearase chain reaction analysis

To measure the expression of the T6SS locus, total RNA was extracted using the TaKaRa MiniBEST Universal RNA Extraction Kit following the manufacturer’s instruction. Reverse-transcription and qRT-PCR was performed according to the Vazyme’s instructions. The *mdh* gene was used as an endogenous control for all the qRT-PCR analyses. The relative transcription levels were calculated using the 2^−ΔΔCt^ method.

### Ethics Statement

The Huashan Institutional Review Board approved the sample collection and will waive the informed consent if further studies are conducted using clinical isolates from the culture collection. Personal privacy is not involved in this study.

## Acknowledgments

We report no potential conflict of interest. This study was supported by the National Natural Science Foundation of China (82173896).

## Data availability

The genomes sequenced during this study are available under the following BioProject accession numbers PRJNA930978. The RNA sequences have been submitted to the NCBI SRA database under BioProject accession number PRJNA972856.

## Supplemental Material

**TABLE S1** Antimicrobial susceptibilities of strains.

**TABLE S2** The expression of T6SS related genes was changed in Δ*hcp* or Δ*vgrG* mutants compared with WT.

**TABLE S3** Strains and plasmids in this study.

**TABLE S4** Primers used in this study.

**FIG S1** T6SS-related gene expression levels in Δ*hcp* or Δ*vgrG* mutants compared with WT by RT-qPCR.

**FIG S2** Interspecies killing by *K. pneumoniae* HS11286 in a contact-dependent manner.

